# Spatial and Single Cell Mapping of Castleman Disease Reveals Key Stromal Cell Types and Cytokine Pathways

**DOI:** 10.1101/2024.09.09.609717

**Authors:** David Smith, Anna Eichinger, Andrew Rech, Julia Wang, Eduardo Esteva, Arta Seyedian, Xiaoxu Yang, Mei Zhang, Dan Martinez, Kai Tan, Minjie Luo, Christopher Park, Boris Reizis, Vinodh Pillai

## Abstract

Castleman disease (CD) is inflammatory lymphoproliferative disorder of unclear etiology. To determine the cellular and molecular basis of CD, we analyzed the spatial proteome of 4,485,009 single cells, transcriptome of 50,117 single nuclei, immune repertoire of 8187 single nuclei, and pathogenic mutations in Unicentric CD, idiopathic Multicentric CD, HHV8-associated MCD, and reactive lymph nodes. CD was characterized by increased non-lymphoid and stromal cells that formed unique microenvironments where they interacted with lymphoid cells. Interaction of activated follicular dendritic cell (FDC) cytoplasmic meshworks with mantle zone B cells was associated with B cell activation and differentiation. VEGF, IL-6, MAPK, and extracellular matrix pathways were elevated in stromal cells of CD. CXCL13+ FDCs, PDGFRA+ T-zone reticular cells (TRC), and ACTA2-positive perivascular reticular cells (PRC) were identified as the predominant source of increased VEGF expression and IL-6 signaling in CD. VEGF expression by FDCs was associated with peri-follicular neovascularization. FDC, TRC and PRC of CD activated JAK-STAT, TGFβ, and MAPK pathways via ligand-receptor interactions involving collagen, integrins, complement components, and VEGF receptors. T, B and plasma cells were polyclonal but showed class-switched and somatically hypermutated IgG1+ plasma cells consistent with stromal cell-driven germinal center activation. In conclusion, our findings show that stromal cell activation and associated B-cell activation and differentiation, neovascularization and stromal remodeling underlie CD and suggest new targets for treatment.

**Key points:**

1. Castleman Disease is characterized by activation and proliferation of CXCL13+ FDCs, PDGFRA+ reticular cells, and ACTA2-positive PRCs.
2. VEGF and IL-6 from lymph node stromal cells are associated with B-cell activation and differentiation, endothelial proliferation, and inflammation in CD

## Introduction

Castleman disease (CD) encompasses a group of disorders characterized by abnormal lymph node morphology(*1–3*). Unicentric hyaline vascular CD (UCD) manifests as the enlargement of one or more lymph nodes in a single anatomical region(*4*), exhibits increased hyalinization and vascularity(*5*) and is typically managed through surgical resection(*6*). Very low level somatic mutational burden has been noted in a subset of UCD(*7*) but the significance and cell of origin in CD is unclear.

Multicentric CD (MCD) is marked by the enlargement of multiple lymph node regions throughout the body and is accompanied by laboratory abnormalities and systemic symptoms(*8*). MCD exhibits a variable lymph node morphology, featuring abnormal germinal centers, increased interfollicular vascularity and plasma cells. MCD is characterized by elevated levels of serum IL-6(*9*), VEGF(*10*), IL-1β and CXCL13(*11*). A subset of MCD cases are associated with HHV-8 infection(*12*) that drives the pathogenesis through viral IL-6(*13*). In contrast, the etiology of HHV8-negative MCD (iMCD) is not known. IL-6 targeted therapy(*14*) and MTOR inhibitors(*15*) are used in the management of iMCD with variable efficacy.

Germline genetic aberrations have been sporadically reported in a subset of iMCD cases(*7, 16, 17*), but the cells initiating disease and contribution to pathogenesis is not clear. Bulk transcriptomics of lymphoid tissue from CD demonstrated that IL-6, VEGF, complement and vascular pathways were upregulated(*18, 19*). Prior immunohistochemical analysis suggested that VEGF and IL-6 expression might originate from lymph node cells(*10, 20*), but conclusive identification of the exact cell type and their role in the pathogenesis of CD was lacking. In our investigation, we aimed to identify the cells and pathways involved in UCD and MCD by utilizing single-cell spatial proteomic and transcriptomic approaches in combination with DNA sequencing and copy number analysis.

## Methods

### Specimen Utilization Workflow

Lymph node resections with clinicopathological diagnoses of UCD, MCD, and non-malignant non-infectious reactive lymph nodes (RLN) were identified from the institutional archives. We identified cases where rapid intraoperative frozen section diagnosis was conducted on whole lymph node resections. Upon removal from the patient, lymph node sections were embedded in Optimal Cutting Temperature (OCT) compound and frozen within a 10-minute window per clinical intraoperative diagnostic requirements that also ensured high-quality RNA reflecting the in-vivo state. The lymph node was bisected along its long axis. One-half of the bisected lymph node was frozen, while the other half was fixed in formalin and subsequently embedded in paraffin (FFPE). The OCT-frozen half was used for single-nuclei transcriptomic analysis, while the corresponding FFPE half was used for single-cell proteomic imaging (Supplemental Figure 1). Frozen tissue was used for DNA sequencing and SNP array analysis.

### Multiplex Spatial Proteomic Imaging and Analysis

#### Image acquisition (CODEX^®^/Phenocycler^®^) and processing

Multiplexed spatial imaging of lymph node cross sections was performed using CODEX^®^/Phenocycler^®^ from Akoya Biosciences. Clinically validated immunohistochemical markers were used to identify major immune and stromal cell types and subsets (Supplemental Table 1). After heat-induced epitope retrieval, a single 5µm thick FFPE tissue was stained with a panel of forty-four DNA oligonucleotide-conjugated marker antibodies per the manufacturer instructions. Sections were then stained with three fluorophores conjugated to DNA oligonucleotides (*21*) and imaged with an inverted microscope (Keyence BZ-X700) using fluorophore-tagged (AF488, Atto550, Cy5) DNA oligonucleotide reporters. The process was iteratively repeated for 19 cycles to image all markers in the panel. Raw images were processed through the CODEX^®^ processing software (CODEX^®^ Processor 1.8.2.13). Different image tiles with similar z-planes were combined to create one qptiff file for analysis. Background subtraction, deconvolution, shading correction and cycle alignment were performed.

#### Lesional region selection and cell segmentation

Aggregated CODEX^®^ Processor images were split into component imaging regions. For each tissue, three to five representative regions were selected for a total of 35 regions. Within these regions, for each tile containing cells, the average channel intensity was collected. Using this sample population, each marker for each tile for all regions was evaluated using a two-tailed Grubb’s test (α = 0.01). If a tile contained more than 8 suboptimal markers/channels, then the tile was excluded from further analysis to ensure high-quality data. Over the entire experiment, this resulted in 211 of 2074 tiles being excluded. Cell segmentation was performed with CellSeg(*22*) using the default model weights. The GROWTH_METHOD and GROWTH_PIXELS parameters were determined through an iterative process, ultimately, set as ‘Sequential’ and 5, respectively. CellSeg was also used for channel intensity quantitation with lateral bleed compensation enabled.

#### Cell phenotyping and identification

To assign cell type labels, data from representative regions of RLN, UCD and MCD (Supplemental Figure 2A) were processed using Scanpy(*23*) v1.3.2. Briefly, marker intensities were log normalized and scaled (Supplemental Figure 2B). Cells were evaluated using Uniform Manifold Approximation and Projection (UMAP) and clustered using the Leiden algorithm(*24*). Marker genes for each cluster were determined using Scanpy’s rank_genes_groups function. Every cluster was assigned a cell type label after expert review of marker expression level and spatial localization of assigned cells (Supplemental Figure 3). These representative data were used to construct a reference data set and predict cell labels for the remaining regions using Symphony(*25*) and Harmony(*26*), respectively. Resultant labels were reviewed and edited in batches through customized marker gating rules. This strategy was successful in annotating the majority of cells. Any remaining indeterminant cells were assigned labels using the phenotyping tools of Scimap (v0.22.9). The phenotyping matrix and gating values for these cells were iteratively determined by expert review. B cells, plasma cells, CD4+ T cells, CD8+ T cells, macrophages, regulatory T cells (Tregs), T follicular helper cells (Tfh), cytotoxic CD8 T cells, classic dendritic cells (cDC) 1 and 2, plasmacytoid dendritic cells (pDC), endothelial cells, lymphatic endothelial cells, neutrophils and mast cells were annotated using this methodology (Supplemental Figure 1). Stromal cells were identified by the lack of pan-hematopoietic marker CD45 or other cell type specific markers in DAPI-positive nuclei. Minor populations of unclassified CD45+ cells were labeled as ‘myeloid’ and unclassified CD3+ cells were labeled ‘immune’.

#### Cell distance analysis and spatial analysis

We used the suite of functionalities from Scimap (www.scimap.xyz) to determine the cellular and spatial architecture. Cell-cell distances and interactions (method=’radius’, radius=50) were calculated on a disease level. For each cell-cell pair combination, the distance distributions were scored using a Kolmogorov–Smirnov test against either the simulated distribution of complete spatial randomness or the empirical values measured in RLN tissue. P-values were adjusted using the Bonferroni method. The η_.50_ distance was used to determine the closest cell pairings in CD compared to RLN.

Spatial neighborhoods were defined using Scimap’s implementation of the Latent Dirichlet Allocation (LDA) method (method=’radius’, radius=50)(*27*). The data were analyzed using 8, 10, 12, and 15 possible microenvironments. Twelve microenvironments provided the optimal balance between known lymph node structures and novel areas. Microenvironments were assigned labels based on unique cell type and spatial distribution.

Image and spatial statistics were calculated for each follicle, as defined by the “B cell, germinal center” and “B cell follicle, stromal enriched” annotations, and averaged for each region. The CD21 area was calculated as the thresholded CD21 fluorescence value and the CD21 signal was calculated as the normalized value in segmented cells. The concept of image entropy was used to quantify the degree of information and complexity in CD21 meshworks. Entropy was calculated as the local entropy within discs of 5-pixel radius. Across a follicle, this is simplified to the root mean square.

### snRNA-seq and analysis

#### Nuclei isolation and sequencing

Frozen tissue was sectioned into 40µm sections and nuclei were recovered using Chromium Nuclei Isolation Kit (PN-1000494) from 10x Genomics. Nuclei integrity and concentration were evaluated using a hemocytometer. Eight thousand nuclei were loaded onto a 10x Genomics Chromium Controller for a targeted recovery of 5,000 nuclei. Single nuclei RNA was processed for sequencing by constructing gene expression (GEX) libraries (Chromium Next GEM Single Cell 3 Kit v3.1, PN-1000268) or gene expression and V(D)J immune profiling libraries (Chromium Next GEM Single Cell 5’ Kit v2, PN-1000263; Chromium Single Cell Human TCR Amplification Kit, PN-1000252; Chromium Single Cell Human BCR Amplification Kit, PN-1000253). Briefly, the single nuclei suspension was mixed with RT Master Mix and loaded with barcoded single-cell 3’ or 5′ gel beads and partitioning oil onto microfluidic chips to encapsulate 5000 nuclei per sample using Chromium Controller. After reverse transcription and cleanup, cDNA libraries were generated according to manufacturer instructions with one additional cycle of PCR amplification to account for the relatively lower amount of RNA in nuclei than in whole cells. cDNA was fragmented and end-repaired, size-selected and PCR amplified to generate a 3’ or 5′ gene expression library. For VDJ library construction, full-length TCR or BCR transcripts were enriched from 4μL of amplified cDNA, and 50 ng of enrichment product was used for library construction. Libraries were submitted for sequencing on an Illumina Novaseq S1-100 flow cell for a minimum sequencing depth of 25,000 reads/nuclei for gene expression and 5,000 reads/nuclei for V(D)J profiling. Reads were processed using CellRanger Single-Cell Software Suite (v6.1.1 and v7.0.0) from 10x Genomics. Reads were aligned to the human reference genome GRCh38. Custom references were also prepared to include Epstein-Barr virus (NCBI:txid10376), human herpesvirus 8 (NCBI:txid37296), and human immunodeficiency virus 1 (NCBI:txid11676).

#### Data integration and annotation

Unique molecular identifier (UMI) counts were corrected for ambient RNA expression in R (v4.1.2) using SoupX(*28*) (v1.6.1). Further processing, visualization, and clustering was performed with Seurat(*29*) (v4.3.0). Briefly, counts were normalized using sctransform, and cells were filtered on a library-specific basis of number of features, total counts, and mtRNA content and rRNA content. Putative doublets were identified using DoubletFinder(*30*) (v2.0.3). Fully processed count data were integrated using Seurat’s RPCA method. After integration and inspection of cell clusters, additional filtering was applied to remove likely cell debris based on feature count. Ultimately, the integrated data set consisted of 50117 nuclei. Ribosomal, mitochondrial, immunoglobulin, and HLA genes were filtered out. Cells were clustered with the Leiden algorithm and cell type labels were manually assigned based on the top marker genes. Subsequent data sets were annotated through label transfer using Symphony(*26*) (v0.1.0).

#### Differential expression and enrichment analyses

Differentially expressed genes (DEG) between RLN, UCD and MCD cases were detected using Wilcoxon Rank sum test implemented with Seurat. GEM libraries generated from 5’ and 3’ sequencing chemistries were batch corrected using ComBat(*31*) (SVA v3.46.0). Cell types were filtered on the criteria that at least 50 cells be represented among test groups. Certain genes were broadly upregulated across multiple cell types. To identify cell-type-specific DEG, the frequency of cluster significance was calculated for each gene, and genes in the top η._96_ were filtered out. Enrichment analysis was conducted with gProfiler2 (v0.2.1). As input to gProfiler, DEG were filtered by p-value<0.1. AUCell(*32*) was used to score the signature of a gene set for each cell.

#### Ligand-receptor interaction analysis

Ligand-receptor interactions between cell-cell pairings were investigated using LIANA(*33*) (v0.1.8). For each case, top scoring interactions were defined by an aggregate score<0.1. Unique interactions of CD compared to RLN were identified. Inter-sample variation was assessed using Tensor-cell2cell(*34*) to identify distinct driver interactions (‘factors’) associated with disease states. Cell types represented in less than 30% of the samples, based on a criterion of 30 observations, were excluded from the analysis. Individual ligand-receptor pair loadings were evaluated for enrichment against a weighted database of pathways provided by PROGENy(*35*).

#### Multiplex RNA *in situ* Hybridization

RNA *in situ* hybridization (ISH) was performed on a Bond Rx (Leica Biosystems) automated staining platform using RNAscope^®^ LS Multiplex assay TSA Vivid dyes (ACD, 323275) according to the manufacturer’s instructions. Probes for *VEGFA* (ACD 423168-C2), *IL-6* (ACD, 310378), *CXCL13*(ACD, 311328-C3), *CXCL12* (ACD, 422998-C3), *PDGFRA* (ACD, 604488-C3), *ACTA2* (ACD, 444778-C3) and *CD19* (ACD, 402718-C3) were obtained. Three RNAscope^®^ probes were stained along with DAPI on sections from cases and controls. Stained slides were digitally scanned at 20X magnification on an Aperio FL slide scanner (Leica Biosystems, Germany). Appropriate positive and negative controls were used per the manufacturer’s recommendation to ensure RNA integrity and exclude background signal.

#### VDJ Sequencing and Analysis

Immune repertoire profiling from V(D)J sequences was performed using the R package, Platypus (v3.5.0). Clonotypes were defined using the “double.and.single.chains” option. For quantitation of somatic hypermutation in B cells, full length sequences were assembled using MiXCR.

#### Targeted DNA sequencing and copy number variant analysis

A targeted next generation sequencing (NGS) panel which interrogates 238 malignancy-associated genes for sequence and copy number variants (CNVs) was performed on the extracted DNA. Briefly, DNA was fragmented and tagged for target enrichment using SureSelect^QXT^ reagent kit (Agilent Technologies to generate adapter-tagged libraries, which were subjected to sequence analysis on the Illumina HiSeq platform for 150 bp paired-end reads (Illumina Inc.). All coding exons and the flanking intron sequences of targeted genes along with selected promoter and intronic regions were sequenced with a targeted average sequence depth of 1800x. NGS data were analyzed using clinical laboratory software ConcordS V2 and NextGENe V2 NGS Analysis Software (SoftGenetics). The annotated sequence variants and CNVs were classified per clinical somatic variant guidelines(*36*).

#### SNP Array Analysis

Genome-wide SNP array analysis was performed on genomic DNA extracted from lymph node tissue of the patients using the Illumina Infinium CytoSNP-850Kv1.2 BeadChip (Illumina Inc). The data were analyzed using vendor-provided analysis software (GenomeStudio). All genomic coordinates were based on the GRCh37/hg19 build of the human genome. The assay detects chromosomal gains, losses, and copy-neutral loss of heterozygosity (cnLOH) involving ≥10 SNP probes that are present in at least 10% of cells.

## Results

### Characteristics of Cases and Controls

The cohort consisted of four MCD, three UCD and two RLN cases (Table 1). The lymph node resections were performed at first presentation of lymphadenopathy. RLN1 with reactive follicular hyperplasia and RLN2 with reactive interfollicular plasmacytosis were controls for the histological features of UCD and MCD lymph nodes. RLN and UCD cases did not show systemic symptoms or laboratory abnormalities except for UCD3. MCD cases comprised three HHV8-negative iMCD (MCD1,2 3), and one HHV8-positive MCD in a patient with HIV (MCD4). MCD4 developed in the setting of HIV infection. iMCD patients presented with fever, anasarca, renal dysfunction, anemia, thrombocytopenia and multicentric lymphadenopathy.

**Table 1.**
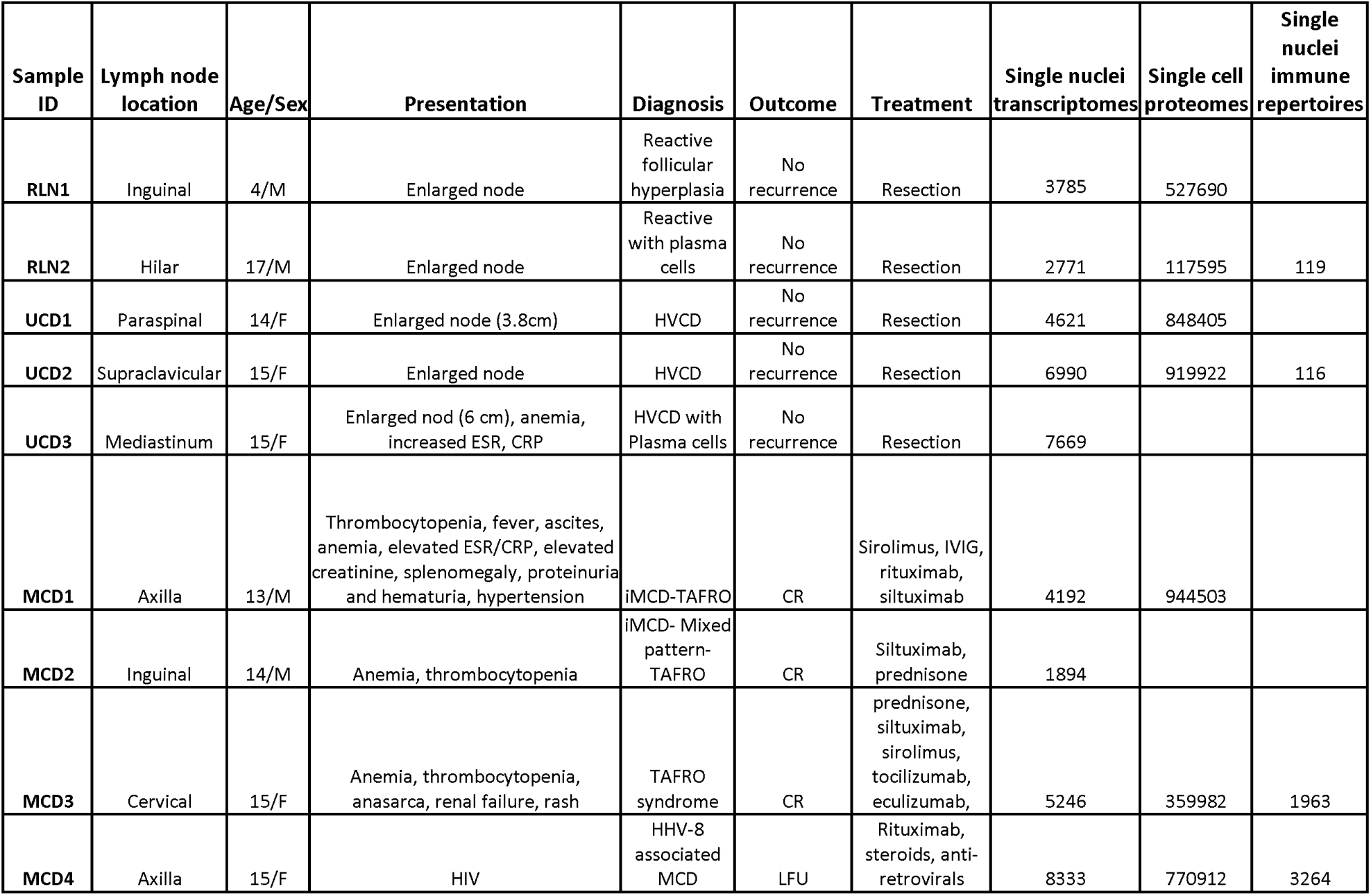
Clinicopathologic Characteristics of Cases and Controls.

### Cellular Composition and Spatial Distribution

The immunophenotype and spatial distribution of 645,285 single cells in RLN, 1,768,327 cells in UCD, and 2,071,397 cells in MCD lymph nodes were analyzed (Figure 1). Cases and controls showed variably sized CD20+ B-cell follicles separated by interfollicular areas with CD3+ T cells, CD11b+ myelomonocytic cells, CD138+ plasma cells, CD31/34+ endothelial cells, and CD45-negative DAPI+ stromal cells (Figure 1A). RLN showed predominantly lymphoid cells (B and T cells) with only minor proportions of the non-lymphoid cell types. Based on permutation tests of MCD vs. RLH, there were significantly (*q* < 0.01 and |log2FC| > 0.53) increased endothelial cells, plasma cells, macrophages, cytotoxic T cells, and stromal cells but decreased B cells compared to RLN (Figure 1B and Supplemental Figure 4). Both UCD and MCD showed decrease in Tfh, Treg and cDC. Strikingly, non-lymphoid cells comprised more than 25% of lymph node cellularity in MCD1 and the majority of lymph node cellularity in MCD3. CD45-negative DAPI-positive stromal nuclei, consistent with follicular dendritic cells (FDCs), were more abundant in the germinal centers of UCD and MCD compared to RLN. However, enumeration of nuclei did not capture the extent and complexity of cytoplasmic projections emanating from FDC nuclei. Cytoplasmic meshworks of FDC play a crucial role in affinity maturation of B cells by presenting native antigen to B cells on complement receptors CD21 (CR2) and CD35 (CR1) (Figure 1C). Hence, we focused on CD21 to visualize the extent of FDC meshworks. Interdigitation of FDC meshworks between concentric layers of B cells led to close interactions and was the basis of the well described ‘onion skin’ appearance of CD mantle zones. Image-analysis techniques were used to quantify the extent and organization of CD21 meshworks. Although the follicles of CD were smaller or similar in size to RLN, the area, brightness, and image entropy (measure of complexity) of CD21 signal was higher in UCD and MCD (Figure 1D). UCD exhibited a lower Shannon diversity index of cells in follicles, consistent with the observed predominance of FDC nuclei. The findings suggested that aberrant activation and proliferation of FDC and other stromal cells may underlie CD.

**Figure 1:**
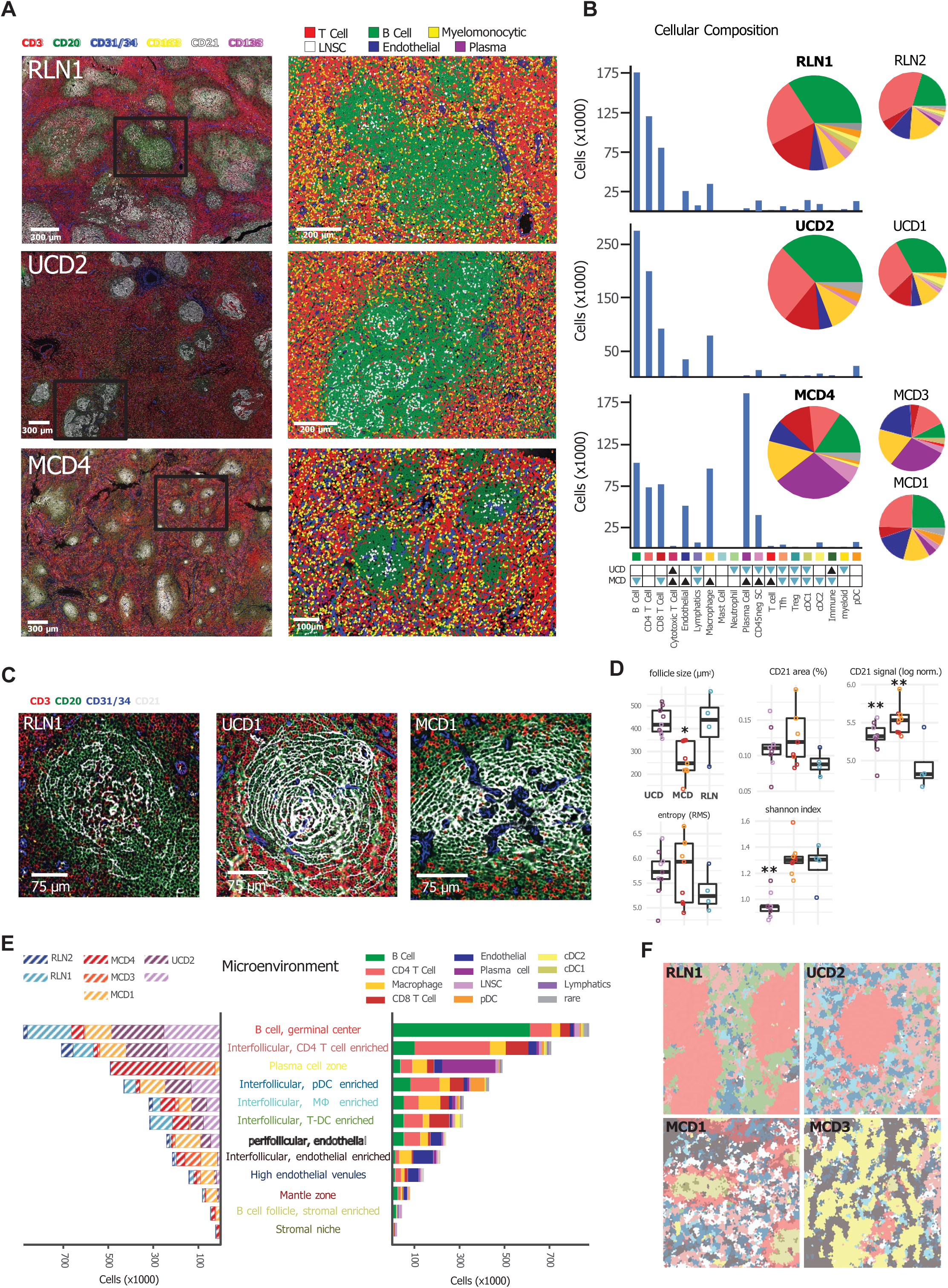
Single-cell proteomic analysis of RLN, UCD and MCD lymph nodes. A. The left column displays representative full scans of RLN1, UCD2, and MCD4, showcasing the expression of key lineage markers. In the right column, cell segmentations and annotations for the indicated germinal center region (black box) are provided. B-cells (green) form follicles, while T cells (red) and myeloid cells (yellow) populate the interfollicular areas. UCD and MCD exhibit increased intrafollicular stromal cell proliferation (white), with MCD additionally showing interfollicular endothelial cell (blue) and plasma cell (magenta) proliferation. RLN, reactive lymph node; UCD, unicentric Castleman disease; MCD, multicentric Castleman disease; CD45negSC, CD45-negative DAPI-positive stromal cells. B. Bar plots illustrate the absolute abundance of cells for the representative region shown in A. Pie charts (inset) represent the relative abundance of cell types across all imaged regions for each sample. Differential abundance against RLN was tested using a permutation test. Significantly (*q* < 0.01 and |log2FC| > 0.53) increased proportions are indicated with black triangles and significantly decreased proportions are indicated with blue triangles. C. Increased CD21+ FDC meshworks (white) in the germinal center regions of UCD and MCD. D. Analysis of size, area, signal intensity, image entropy of CD21, and cell diversity (Shannon index) within each follicle reveals higher expression and organization of FDC meshworks across CD, and low cell diversity in UCD. Significance versus RLN1 was calculated using Tukey’s Honest Significance Difference test. *, *p* < 0.05; **, *p* < 0.01. E. Twelve identified microenvironments are depicted. Sample composition (left graph) and cellular composition (right graph) of each microenvironment are shown. B cell germinal center and interfollicular CD4-and T-DC -enriched microenvironments are present in all samples. Plasma cell, macrophage, endothelial and stromal enriched microenvironments are increased in CD samples. F. Voronoi diagrams of microenvironments in representative regions of cases and controls are shown. Color scheme as indicated by shading of microenvironment names in 1E.

### Microenvironments and Cellular Interactions of CD

Next, we assessed the differences in the microenvironment between cases and controls. Twelve microenvironments were identified from the combined spatial scans of RLN, UCD, and MCD regions (Figure 1E and F). B-cell germinal centers and interfollicular microenvironments enriched in CD4 T cells, macrophages or DC were noted, consistent with known lymph node structures. Many microenvironments were enriched in CD cases. An interfollicular macrophage-enriched microenvironment was noted in all CD suggestive of increased macrophage activity. Two endothelial-predominant microenvironments (perifollicular and interfollicular) and a plasma-cell region were identified in all MCD. UCD1/2, and MCD1 exhibited unique perifollicular-endothelial and plasmacytoid dendritic cell (pDC) regions, while MCD3 and MCD4 showed a predominance of interfollicular-endothelial and plasma cell microenvironments. MCD1 showed a unique mantle microenvironment and an stromal cell-rich B-cell follicle was present in MCD1 and MCD4. The microenvironments comprised multiple other cell types that likely interacted with each other akin to the functional interactions that result in normal lymph node structures. We hypothesized that interactions between locally proximal cells may drive the pathogenesis of CD. The distribution of distances between different cell types in each sample was calculated and tested against their RLN counterparts using a Kolmogorov Smirnov test (Supplemental Table 2). Cell-cell pairings with an average distance <50μm were considered biologically relevant. The cell-cell interactions in UCD and MCD that were most different from RLN were those of plasma cells, macrophages, endothelial cells with other cell types. These findings suggest that the distinctive microenvironments wherein non-lymphoid cells engage with lymphoid cells play a role in the pathogenesis of CD.

### Functional analysis of expanded cell populations in CD

Single cell proteomics provided accurate enumeration of major cell types and microenvironments of CD but provided limited information on cell function. Hence, we performed single-nuclei RNA-seq of concurrent frozen tissue to characterize cell states and signaling pathways. Single-nuclei transcriptomes for 6,556 cells from RLN, 19,280 from UCD and 24,281 from MCD lymph nodes were analyzed (Figure 2). Distinct populations of B-cell, plasma-cell, T-cell, myelomonocytic and stromal lineages were identified (Figure 2A). RLN showed abundant germinal center B-cell nuclei that were decreased in UCD and MCD. Based on permutations tests against RLN, UCD showed significantly (*q* < 0.01 and |log2FC| > 0.53) increased pDC, plasma cells and naïve B cells. MCD showed increased nuclei of plasma cells, proliferating plasma cells (plasmablasts), endothelial cells, lymphatics, fibroblastic reticular cells (FRC), monocytes and macrophages and cytotoxic T cells. Although non-lymphoid populations were underrepresented in transcriptomic data (likely due to dissociation limitations), the relative composition of expanded cell nuclei closely resembled the cell types observed in proteomic data.

**Figure 2:**
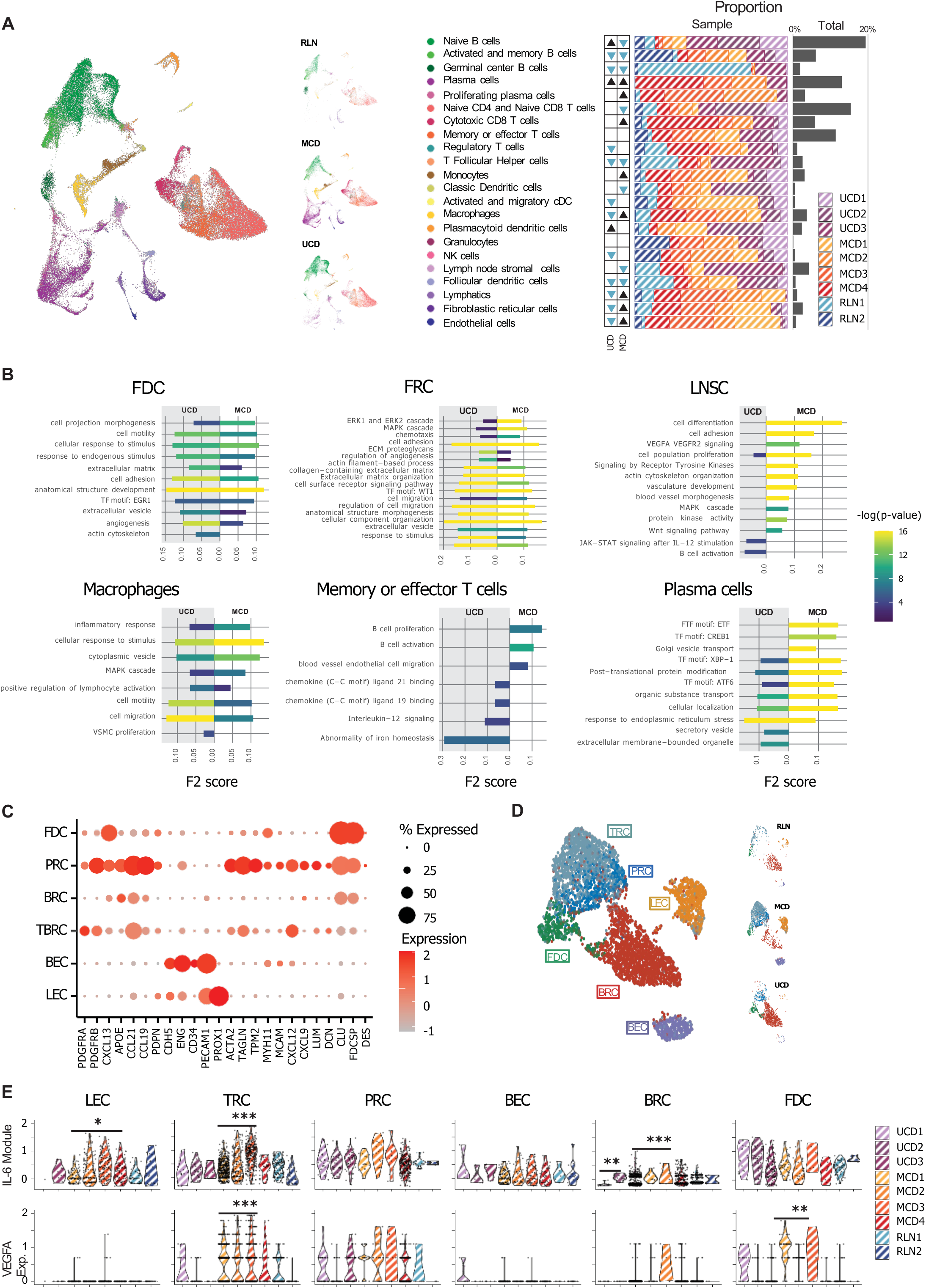
Comparative Analysis of Single Nuclei RNA-Sequencing and Pathway Enrichment in Cell Types of UCD and MCD versus RLN. A. A UMAP plot of 50,117 single nuclei displays identified cell types and their relative abundance per sample, along with global abundance. MCD exhibits significantly increased proliferating plasma cells, cytotoxic and memory T cells, monocyte-macrophages, endothelial-lymphatics, and fibroblastic reticular cells (FRC). UCD demonstrates significantly increased plasmacytoid dendritic cells, plasma cells and naïve B cells. Differential abundance against RLN was tested using a permutation test. Significantly increased proportions are indicated with black triangles and significantly decreased proportions are indicated with blue triangles. B. The differentially expressed genes (DEGs) of UCD and MCD compared to RLN were tested for pathway enrichment using various databases, including Reactome, GO, KEGG, and TRANSFAC. The top significantly enriched pathways for MCD and UCD are displayed. LNSC showed enrichment in pathways related to the extracellular matrix (ECM), cytoplasmic projections, actin cytoskeleton, angiogenesis, MAPK signaling, and VEGF signaling. Macrophages are enriched in inflammatory pathways, whereas plasma cells are enriched in pathways associated with antibody production. C. Stromal subsets were annotated by expression of key lineage defining markers. FDCs are characterized by strong expression of *CXCL13*, *CLU* and *FDSP*. PRCs show high expression level of *PDGFRB*, *CCL21*, *CCL19*, *ACTA2*, *TAGLN* and *CXCL12*. TRCs show high *CCL21* and *CXCL12* expression. BECs show high *CDH5*, ENG, and *PECAM1,* while LECs show high *PROX1*. D. UMAPs of merged stromal subsets from cases and controls. TRCs, PRCs, BECs and LECs are predominantly from MCD. BRCs and FDCs are predominantly from UCD. E. SCT normalized IL-6 module score (*IL-6*, *OSMR*, *IL-6ST*, *LIFR*) expression is displayed in the top row while *VEGFA* categorized by stromal cell and sample type is shown in the bottom row. CRCs, PRCs and FDCs of CD show high IL-6 module and VEGF expression. Gene expression significance versus RLN was calculated using Wilcoxon signed-rank test and module score significance versus RLN was calculated using Tukey Honest Significance Difference. *, *p* < 0.05; **, *p* < 0.01; ***, *p* < 0.001.

We next sought to define functional differences of each cell type between cases and controls. We performed differential gene expression and pathway enrichment analyses of UCD and MCD cell types compared to those in RLN (Figure 2B). In UCD and MCD, FDCs exhibited an upregulation of pathways linked to cytoplasmic projections and angiogenesis. Stromal populations displayed an elevated expression of MAPK, angiogenesis, and extracellular matrix-associated pathways. Notably, FRC of MCD showed a significant (p<0.05) increase in VEGF, angiogenesis, MAPK, and JAK-STAT pathways. Macrophages were enriched in inflammatory pathways, while memory T cells were enriched in pathways associated with CCL19/21 binding, B-cell activation, and proliferation. IL-6 pathways were upregulated in activated B cells. Plasma cells were enriched in pathways indicative of increased antibody production. Collectively, these findings indicate that CD is marked by the involvement of various cell types and processes, including increased angiogenesis and extracellular matrix remodeling by stromal populations, inflammation by macrophages, T-cell activation, and IL-6-mediated differentiation of B cells into antibody-producing plasma cells. While these processes are part of normal immune responses, they appear to be hyperactive in CD.

### Stromal Cells and Origin of Key Cytokines

Since key pathways were enriched in stromal cell populations of CD, we analyzed them in greater detail (Figure 2C-E). Single-nuclei transcriptomes from all non-hematopoietic cell types in all samples were extracted and annotated using stromal-specific markers (Figure 2C). We identified distinct groups such as *CDH5*+ *ENG*+ blood endothelial cells (BECs) and *PROX1*+ lymphatic endothelial cells (LECs), which were particularly prevalent in MCD samples. We also noted a predominance of *ACTA2*+ perivascular reticular cells (PRCs) and *PDGFRA/B*+ *CCL21*high *CCL19*low *CXCL12*+ T-zone reticular cells (TRCs) in MCD. In contrast, *CXCL13*+ FDC and *FDCSP*+*CLU*+ B-zone reticular cells (BRCs) were mainly found in UCD samples (Figure 2D). The findings suggest that distinct populations of stromal subsets are involved in the pathogenesis of UCD and MCD.

Considering the elevated levels of circulating VEGF and IL-6 typically seen in CD, we compared their expression across different samples and cell types (Figure 2E). The highest expression of *VEGFA* and IL-6-associated genes (IL-6 module) were observed in stromal populations. Notably, high levels of *VEGFA* were uniquely seen in PRCs and TRCs across all MCD samples (Figure 2E, top row). *VEGFA* was also expressed in FDCs across various CD samples, specifically UCD1, MCD1, and MCD3. Similarly, high expression levels of IL-6 associated genes were observed in PRC, CRC of MCD, and FDCs of UCD (Figure 2E, bottom row). These results showed that stromal cells were the primary source of significant cytokines in CD.

### Spatial Localization of VEGF and IL-6 Expressing Stromal Cells

A sensitive multiplex nucleic acid in-situ hybridization (ISH) was used to co-localize key cytokines and stromal cells. RNAscope assays were performed for *VEGFA*, *IL-6*, *CD19*, *CXCL12*, *CXCL13*, *PDGFRA*, and *ACTA2* (Figure 3A). *VEGFA* exhibited high expression levels in both UCD and MCD, albeit with distinct patterns. In UCD, *VEGF* expression was confined to the follicles, whereas MCD showed markedly elevated expression in both follicular and interfollicular regions (Figure 3A). Within the follicles of both UCD and MCD, *VEGF* was strikingly expressed in the nuclei of FDCs with cytoplasmic *CXCL13* (Figure 3A, left column). In the interfollicular space, *VEGF* co-localized with *PDGFRA*-expressing stromal cells that were consistent with TRCs (Figure 3A, middle column). Additionally, *VEGF* co-expressed with *ACTA2*-expressing PRCs that were associated with blood vessels. IL-6 demonstrated strong expression in MCD3 and 4 (Figure 3A, right column). MCD3 showed co-expression of IL-6, VEGF and PDGFR while MCD4 showed co-expression of IL6 and PDGFR. Co-expression of IL-6 with *CD19*+ immunoblasts and ACTA2+ PRCs was also observed in MCD4.

**Figure 3.**
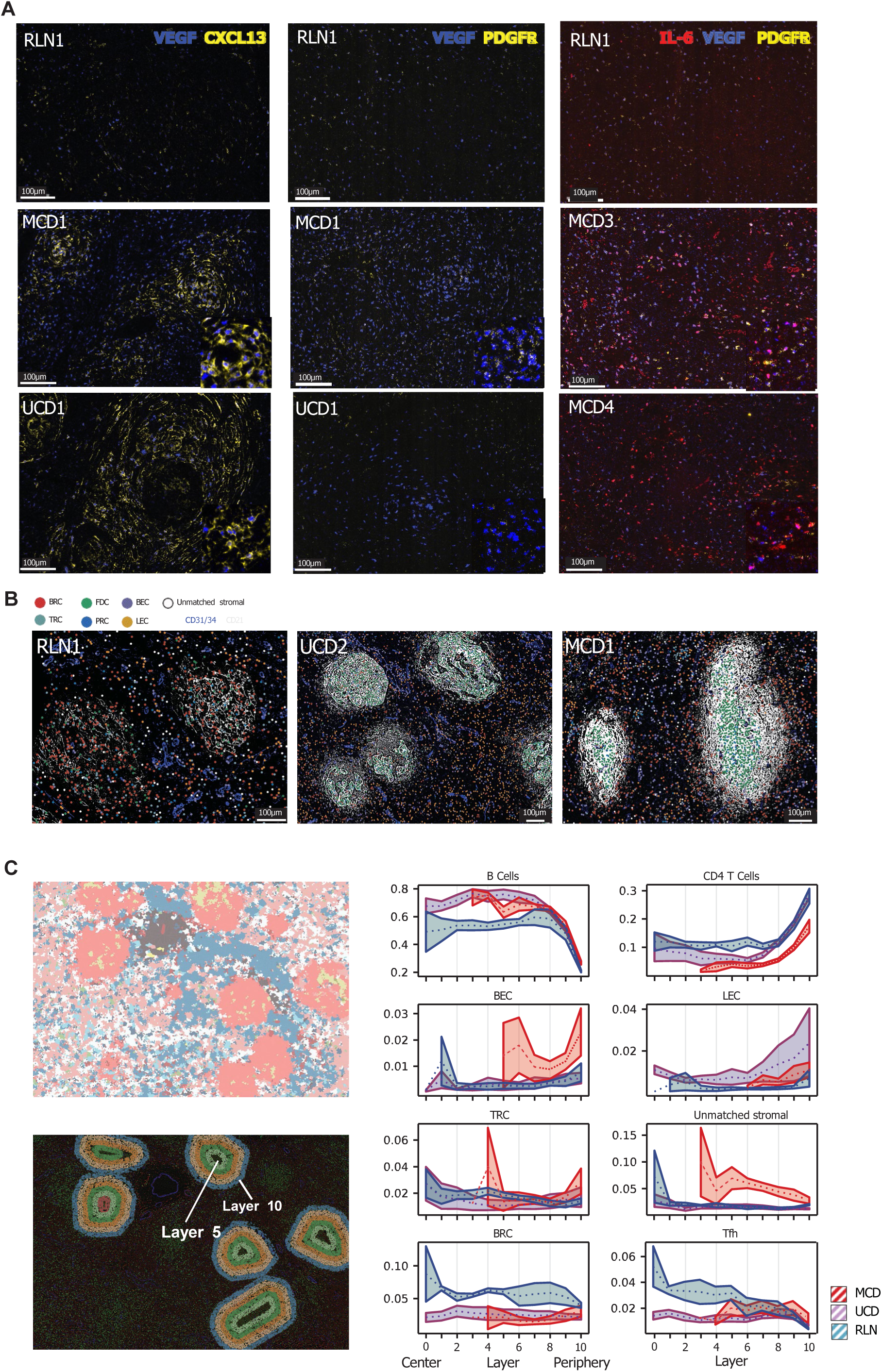
Spatial Localization of Stromal Subtypes. A. RNA ISH for *VEGFA/CXCL13*, *VEGF/PDGFRA* and *IL-6/VEGFA/PDGFRA* combinations in representative regions of RLN, UCD and MCD are shown. Markedly increased *VEGF* and *IL-6* in MCD that colocalized with *CXCL13*+ FDC in follicles and *PDGFRA*+ stromal cells in interfollicular areas (2^nd^ row). Colocalization of *CXCL13*+ FDC and *VEGF* expression in follicles of UCD is noted (3^rd^ row). Co-expression of IL-6, VEGF and PDGFR+ stromal cells in MCD3 is shown (3^rd^ column, purple cells). Co-expression of IL-6 with PDGFR+ stromal cells in MCD4. Inset image shows high magnification. B. Stromal subsets identified from snRNA-seq data are shown in CODEX images after integration by MaxFuse. BEC, LEC and PRC are associated with CD31/CD34+ blood vessels and are increased in UCD and MCD. Markedly increased FDC nuclei are closely associated with regions of dense CD21 meshworks. C. Left column: Follicles were identified by cell microenvironment (Voronoi plot shown in top) and split into concentric 100-pixel contours (bottom). Stromal subtypes identified from snRNA-seq data were transferred to multiplexed image data and their abundance assessed at each contour layer (middle and right columns). Peaks of stromal cells are evident in layers before peaks of BEC and LEC of MCD. MCD is shown in red, UCD in purple and RLN in blue.

### Integration of transcriptomic and proteomic data

Multiple stromal subsets were identified through SnRNA-seq, but accurate distinction on CODEX data was challenging due to the lack of specific surface markers. To address this, we integrated transcriptomics and proteomics data using MaxFuse(*37*) to determine the spatial localization of stromal subsets. This integration allowed us to identify the spatial distribution of key stromal subsets in and around germinal center follicles (Figure 3B). FDC nuclei were accurately localized to the center of CD21-positive meshworks and were markedly increased in UCD and MCD. BRC were localized within follicles of RLN and UCD. PRC were detected in the blood vessel walls and were distinct from BEC and LEC.

Since key cells and pathways of CD were associated with germinal center follicles, we performed a detailed analysis of their cellular composition. Follicles identified by cell-microenvironment analysis were separated into concentric 100-pixel contours (Figure 3C, left column). Stromal subtypes identified from snRNA-seq data were transferred to multiplexed image data and their abundance assessed at each contour layer. B cells and CD4+ T cells were increased in follicular and perifollicular areas respectively. BEC showed a peak in the perifollicular region that was adjacent to peaks of stromal cells in MCD. The spatial findings are consistent with tropism of endothelial cells towards VEGF secreted by stromal cells and formed the basis of the diagnostic feature of penetrating blood vessels (‘lollipop’) in CD.

### Ligand-Receptor Interactions of Stromal Cells

Ligand-receptor interactions that drive UCD and MCD (Figure 4 and Supplemental Figure 6) were identified from snRNA-seq. LIANA(*38*) in combination with Tensor-cell2cell(*34*) identified several factors that distinguish disease types (Figure 4A). UCD was characterized by factors representing stromal cell interactions that activate the JAK-STAT, TGFβ, and MAPK pathways. The ligand-receptor pairs identified in UCD could be categorized as collagen-integrin extra cellular matrix (ECM) interactions, complement-mediated inflammatory interactions, and VEGF interactions between PRC and macrophages(Figure 4B and Supplemental Figure 5).

**Figure 4.**
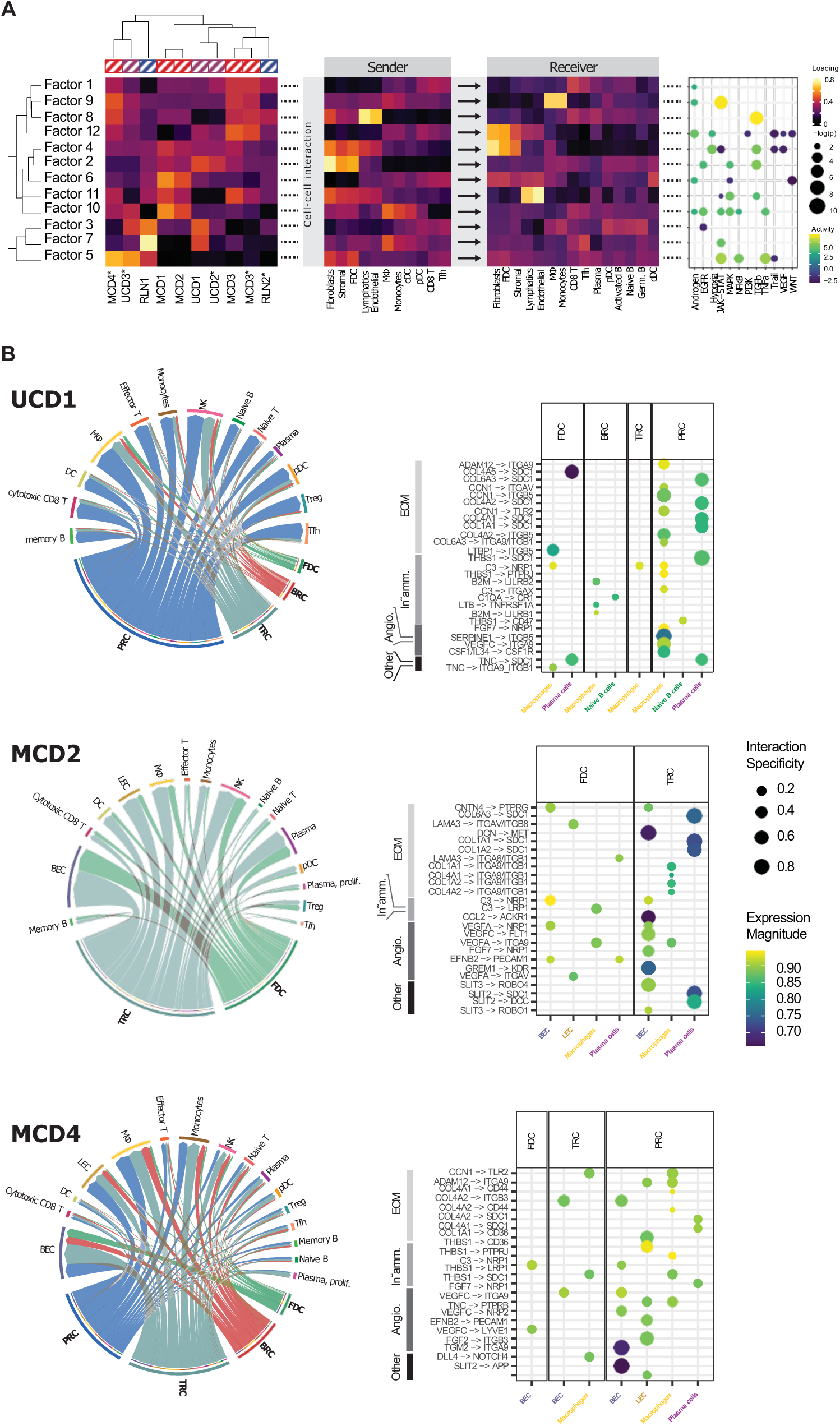
Driving Factors and Ligand-Receptor Interactions in CD. A. Ligand-receptor interactions were assessed for each sample using the LIANA workflow. Inter-sample variation was analyzed with Tensor-cell2cell to identify distinct driver interactions (‘factors’) associated with disease states. Heatmaps display sample identity, source cell type, and receiver cell type for each factor. Enriched pathways for each factor are shown in the dot plot. HVCD1 and 2 are characterized by factor 2. MCD1, 2 and 3 are characterized by factors 6, 10, and 11. MCD3 and 4 are additionally characterized by factors 1, 8, 9, and 12. *denotes samples profiled using 5’ chemistry. B. Significant (p < 0.05) LIANA-identified interactions of UCD and MCD cases compared to RLN are shown. Circos plots and dot plots with stromal subtypes as senders and all other cell types as receivers are shown. Dot plots illustrate individual ligand-receptor interactions clustered based on the Gene Ontology (GO) of the ligand. Interactions are grouped into categories representing ECM (integrin/collagen), inflammatory (complement components), and angiogenesis (VEGF) interactions.

MCD was enriched in factors that reflect the interactions of stromal cells, endothelial cells, macrophages, CD8 T cells and Tfh resulting in the activation of JAK-STAT, TGFβ, MAPK and TNFα pathways. The ligand-receptor pairs identified in MCD2 were interactions of TRCs with plasma cells through collagens and SDC1/CD138 (Figure 4C and Supplemental Figure 5). TRCs showed interactions of VEGF with its receptors on BEC. MCD4 exhibited similar interactions of PRC with BECs and LECs. These findings reveal key ligand-receptor interactions that could be targeted to alleviate MCD symptoms.

### DNA sequencing, Immune Repertoire, and Viral Sequence Analysis

Clinical NGS and SNP array DNA analysis did not reveal clinically significant Tier 1 or 2 single nucleotide or copy number variants (Supplemental Table 3). Several Tier 3 variants of uncertain significance with high variant allele frequencies (VAFs) were noted in UCD1,3 and MCD1,4 suggesting possible germline variants. Reanalysis of data for variants previously reported in CD(*7*) revealed very low levels of *PDGFRB* c.1997 A>G, p.N666S (UCD2), *ALK* c.875G>A, p.R292H (UCD2,3, MCD1,3) and *BCOR* c.3866G>A, p.G1289D (UCD3) in some cases of UCD and MCD. These rare sequencing reads (2-5/1000) were close to the error rate of Illumina sequencing (1/1000) and thus are best considered as variants of uncertain significance.

5’ sequencing was employed to analyze the T and B cell immune repertoire in a subset of cases. 1,631 single nuclei TCR and 6,556 single nuclei BCR sequences were identified from RLN, UCD and MCD. Clonally restricted populations of B or T cells were not observed. VDJ and VJ analyses revealed that plasma cells of UCD3 and MCD3/4 showed high levels of somatic hypermutation and were predominantly class switched to IgG1 (Supplemental Figure 6A and B) suggesting generation from a follicular reticular cell-driven germinal center process.

Given the association of MCD with HHV8 and HIV infection, we aligned sequencing reads to HHV8, HIV and EBV reference genomes. HHV8 sequences were only identified in MCD4 with the highest proportion of viral sequences in plasma cells (Supplemental Figure 6C). The findings were consistent with the HHV8-driven etiology in MCD4. HHV-8, EBV or HIV sequences were not detected in any other UCD, MCD or RLN.

In conclusion, the findings indicate that the activation and proliferation of follicular reticular cells, resulting in VEGF and IL-6 secretion, are linked to B-cell activation, plasma cell differentiation, angiogenesis, and stromal remodeling across all subtypes of CD.

## Discussion

Castleman Disease refers to a heterogeneous group of disorders characterized by pathognomonic lymph node features. CD lymph nodes show similar histological features of germinal center depletion, plasmacytosis, increased vascularity and hyalinization but the mechanistic basis of these features and their treatment implications are unclear. The investigation of CD has been constrained by its unpredictable presentation, and lack of cell culture or murine models. To address these limitations, our study leveraged a rare archive of fresh frozen and FFPE samples obtained from whole lymph node resections. We performed the largest spatial and single cell characterization of lymph node samples and identify novel pathogenic cell types and pathways of pathogenesis in CD. Our findings highlight the immune basis of CD and suggest novel approaches for treatment. The findings will also enable the development of accurate in-vitro and in-vivo models of CD.

We showed that UCD is characterized by activation and proliferation of specific stromal cells resulting that are responsible for the diagnostic morphologic and clinical features. The classic ’onion skin’ appearance of UCD is due to proliferating FDC meshworks interdigitating concentrically between B cells. Another key diagnostic feature of CD ‘twinning/shared mantle zones’ could be attributed to robust proliferation and fusion of separate FDC and their meshworks. CXCL13 and CXCL12-expressing follicular reticular cells are crucial for germinal center activation(*39*). In CD, this activation can result in either proliferating or depleted germinal centers, depending on the time elapsed since the initiating event. We showed that IL-6 and VEGFA were highly expressed by follicular reticular cells. IL-6 is a key cytokine that drives germinal center B-cell activation and plasma-cell differentiation(*40*). In CD, this results in proliferation of plasma cells. High VEGF expression from FDCs is likely basis of characteristic ‘lollipop sign’-blood vessels penetrating FDC-predominant follicles. Functionally, VEGF-driven endothelial proliferation and macrophage driven stromal remodeling is necessary for lymph node enlargement during immune response(*41*). The overall findings suggest that lymphadenopathy in CD may be an immune reaction to aberrant FDC activation.

Similar to UCD, MCD also showed a proliferation and activation of lymph node stromal cells. MCD is characterized by an aberrant immune response with elevated levels of CXCL13, IL-6 and VEGF. The population underlying increased VEGF signaling in MCD were TRCs, PRCs and FDCs. VEGF from stromal cells interacted with its receptors on BECs and LECs to stimulate angiogenesis. The resulting vascular leak from abnormally proliferating blood vessels likely underlies edema, ascites and effusions in MCD. MCD also shows features of germinal center proliferation and plasma-cell generation. Normally, FDC plays an important role in the selection of high-affinity B cells through sustained presentation of native antigen on complement receptors(*42*). FDC-secretion of IL-6 drives the differentiation of B cells to antibody-producing plasma cells(*40*). MCD may represent aberrant activation of follicular reticular cells resulting in expansion of pathogenic antibody-producing plasma cells that result in gammaglobulinemia.

Our findings a cellular and molecular classification of CD with treatment implications. Since FDC play a central role in the pathogenesis of CD, targeting their activation may be an alternate strategy to mitigate IL6, VEGF and CXCL13 cytokinemia. Lymphotoxin B receptor fusion proteins that block FDC activation (*43*) needs further investigation in CD. Presence of CD20-expressing proliferating plasmablasts in plasma cell-predominant CD suggests that CD20-targeting agents may more useful in some case of CD. While rituximab as a monotherapy has limited efficacy in MCD, combination therapies involving CD20 monoclonal antibodies with IL-6 inhibitors or proteasome inhibitors in the plasma cell subtype of iMCD needs further investigation. Systemic and local VEGF inhibitors may be considered in endothelial-predominant subtypes of MCD and UCD.

## Acknowledgements

We would like to thank Ms. Rachel Olson, Mr. Brian Lockhart, and the Division of Hematopathology for their support.

## Ethics Approval

The study was approved by the CHOP Institutional Review Board (IRB 16-013199) and performed in accordance with the Declaration of Helsinki.

## Author contributions

V.P. performed study concept and design; V.P., D.S., A.E., J.W., A.R., K.T., M.Z., M.L., C.P. and B.R., performed writing, review, and revision of the paper; V.P., D.S., A.E., A.S., X.Y., D.M., J.W., A.R. and M.L. provided acquisition, analysis and interpretation of data, and statistical analysis; All authors read and approved the final paper.

## Data availability statement

Data generated or analyzed during this study are included in this published article (and its supplementary information files). Any other data will be provided on reasonable request to the corresponding author.

