## Supplemental Data for "Spatial and Single Cell Mapping of Castleman Disease Reveals Key Stromal Cell Types and Cytokine Pathways"

**Lymph Node Stromal Cells Underlie the Clinicopathologic Features of Castleman Disease**

**Short title:** Cellular and Molecular Basis of Castleman disease

**Key points:**

1. UCD and MCD are characterized by activation and proliferation of lymph node stromal populations.
2. VEGF and IL-6 from lymph node stromal cells drive B cell activation and differentiation into plasma cells, endothelial cell proliferation, macrophage mediated inflammation, and stromal remodeling in CD

Corresponding author:

Vinodh Pillai., MD, PhD. Associate Professor of Pathology and Laboratory Medicine, University of Pennsylvania. Electronic address:.

**Supplemental Material**

**Supplemental Table 1. Categorization of immunophenotypic markers used in CODEX.**

**Supplemental Table 2. Cell-Cell distance analysis.** Closest cell-cell distances in MCD and UCD ordered by difference from RLN. The distribution of distances between different cell types in each sample was calculated and tested against their RLN counterparts using a Kolmogorov Smirnov test (Supplemental Table 2). Cell-cell pairings with an average distance <50μm were considered biologically relevant.

**Supplemental Table 3. Genomic analysis of UCD and MCD.** Copy number variants (CNV) and sequence variants including single nucleotide variants (SNV) and small Insertions (Ins) and Deletions (Del) identified through SNP array and targeted DNA sequencing analyses. The variant allele frequency for each variant is shown within parentheses.

**Supplemental Figure 1. Specimen utilization workflow.** Parallel sections of FFPE and OCT-embedded frozen tissue were used for single cell proteomic, single nuclei transcriptomic and genetic analysis.

**Supplemental Figure 2. A. Key cell markers and immunophenotypic heatmaps in representative regions of RLN1, UCD2, MCD1 and MCD3.** CD3+ T cells are depicted in red, CD20+ B cells in green, CD31/34+ vasculature in blue, CD138+ plasma cells in magenta and CD163+ histiocytes in yellow. B. Heatmap of marker expression by cell type in the representative region of RLN1, UCD2, MCD1 and MCD3 shown in A.

**Supplemental figure 3. Expression of key cell markers and annotations of major cell types.** Cell marker expression is depicted in red in while cell annotations are depicted in green. All nuclei are highlighted by DAPI in white. A. CD3 expression and T cell annotation. B. CD20 expression and B cell annotation. C. CD31 and PDPN expression, endothelial and lymphatic annotation. D. CD11b expression and myelomonocytic cell annotation. E. CD123 expression and plasmacytoid dendritic cell annotation. F. DAPI-positive and CD45-negative stromal cell annotation. G.CD1c expression and dendritic cell annotation.

**Supplemental Figure 4.** **Aggregated single cell proteomic characterization of cases and controls.**  A. Bar plots display the average number of cells across all imaged regions of RLN, UCD and MCD. The inset pie chart illustrates the relative composition of cells aggregated across imaged regions of RLN, MCD and UCD. MCD is characterized by increased plasma cells, macrophages, endothelial cells and stromal cells. B. Differential abundance of MCD and UCD cell populations tested against RLN by permutation test. Significantly increased populations (left) and significantly downregulated populations (right) are shown. by increased plasma cells, macrophages, endothelial cells and stromal cells.

**Supplemental Figure 5.** Ligand receptor interactions in UCD2, UCD3 and MCD2 and MCD3. UCD and MCD are characterized by VEGF, complement component, integrin and collagen interactions.

**Supplemental Figure 6.** A. High levels of somatic hypermutation in plasma cells of UCD3, MCD3 and MCD4. B. Plasma cells are predominantly class switched IgG1 isotype. C. HHV8 viral sequences detected in many cell types of MCD4. Highest levels of HHV8 in plasma cells and cytotoxic CD8+ T cells.
