## Supplemental Tables for "Spatial and Single Cell Mapping of Castleman Disease Reveals Key Stromal Cell Types and Cytokine Pathways"

### Slide 1
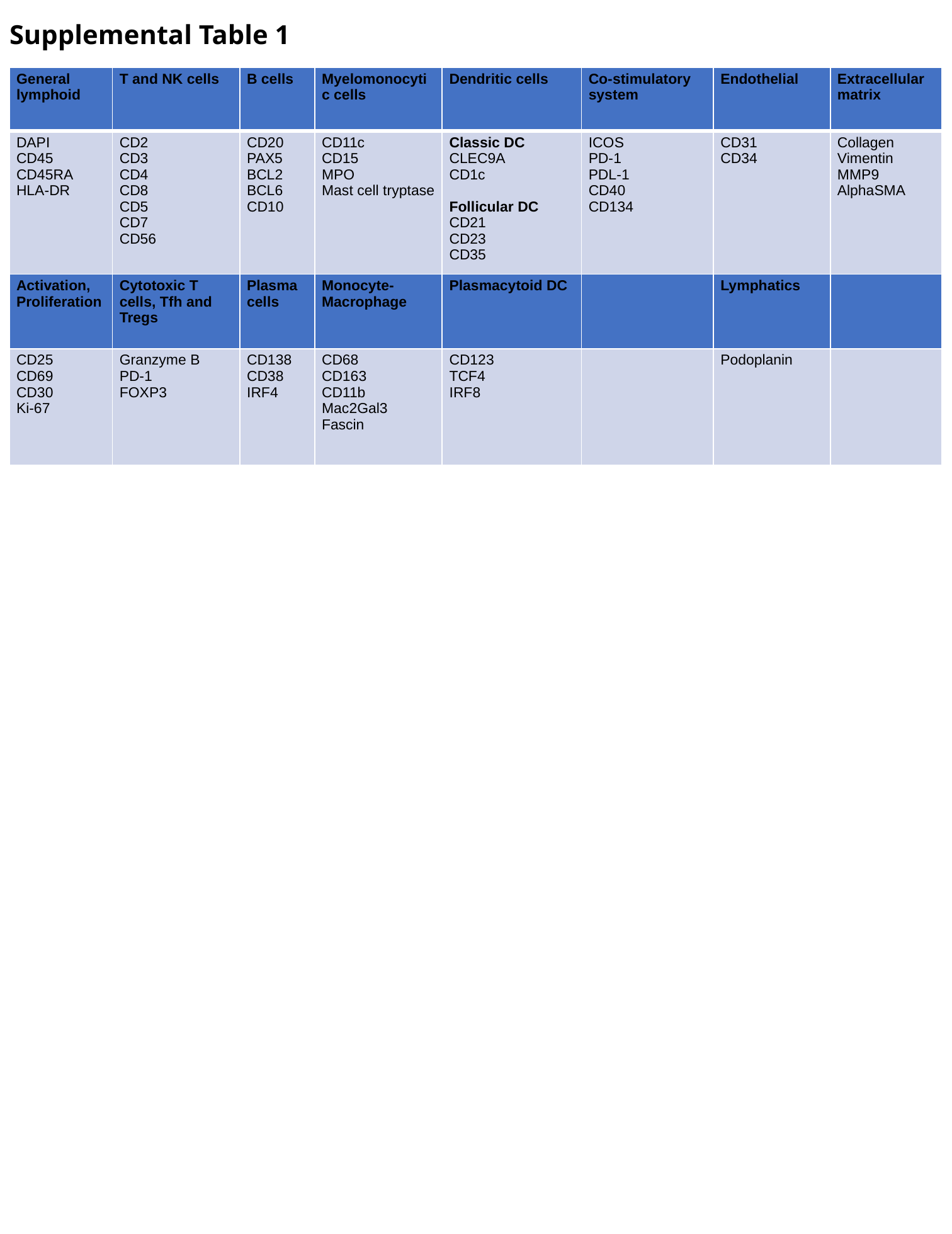

Supplemental Table 1
| General lymphoid | T and NK cells | B cells | Myelomonocytic cells | Dendritic cells | Co-stimulatory system | Endothelial | Extracellular matrix |
| --- | --- | --- | --- | --- | --- | --- | --- |
| DAPI CD45 CD45RA HLA-DR | CD2 CD3 CD4 CD8 CD5 CD7 CD56 | CD20 PAX5 BCL2 BCL6 CD10 | CD11c CD15 MPO Mast cell tryptase | Classic DC CLEC9A CD1c Follicular DC CD21 CD23 CD35 | ICOS PD-1 PDL-1 CD40 CD134 | CD31 CD34 | Collagen Vimentin MMP9 AlphaSMA |
| Activation, Proliferation | Cytotoxic T cells, Tfh and Tregs | Plasma cells | Monocyte-Macrophage | Plasmacytoid DC | | Lymphatics | |
| CD25 CD69 CD30 Ki-67 | Granzyme B PD-1 FOXP3 | CD138 CD38 IRF4 | CD68 CD163 CD11b Mac2Gal3 Fascin | CD123 TCF4 IRF8 | | Podoplanin | |

### Slide 2
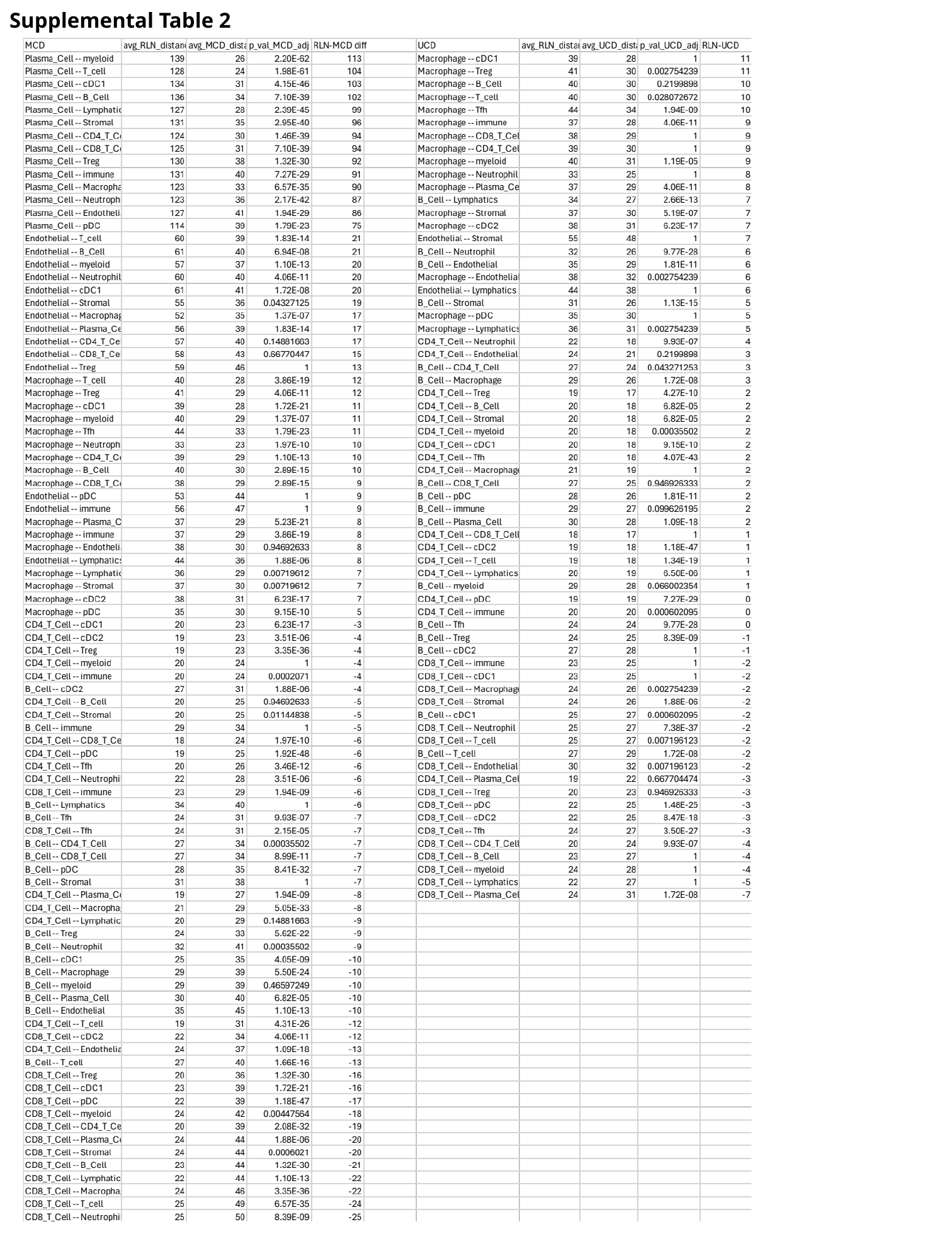

Supplemental Table 2

### Slide 3
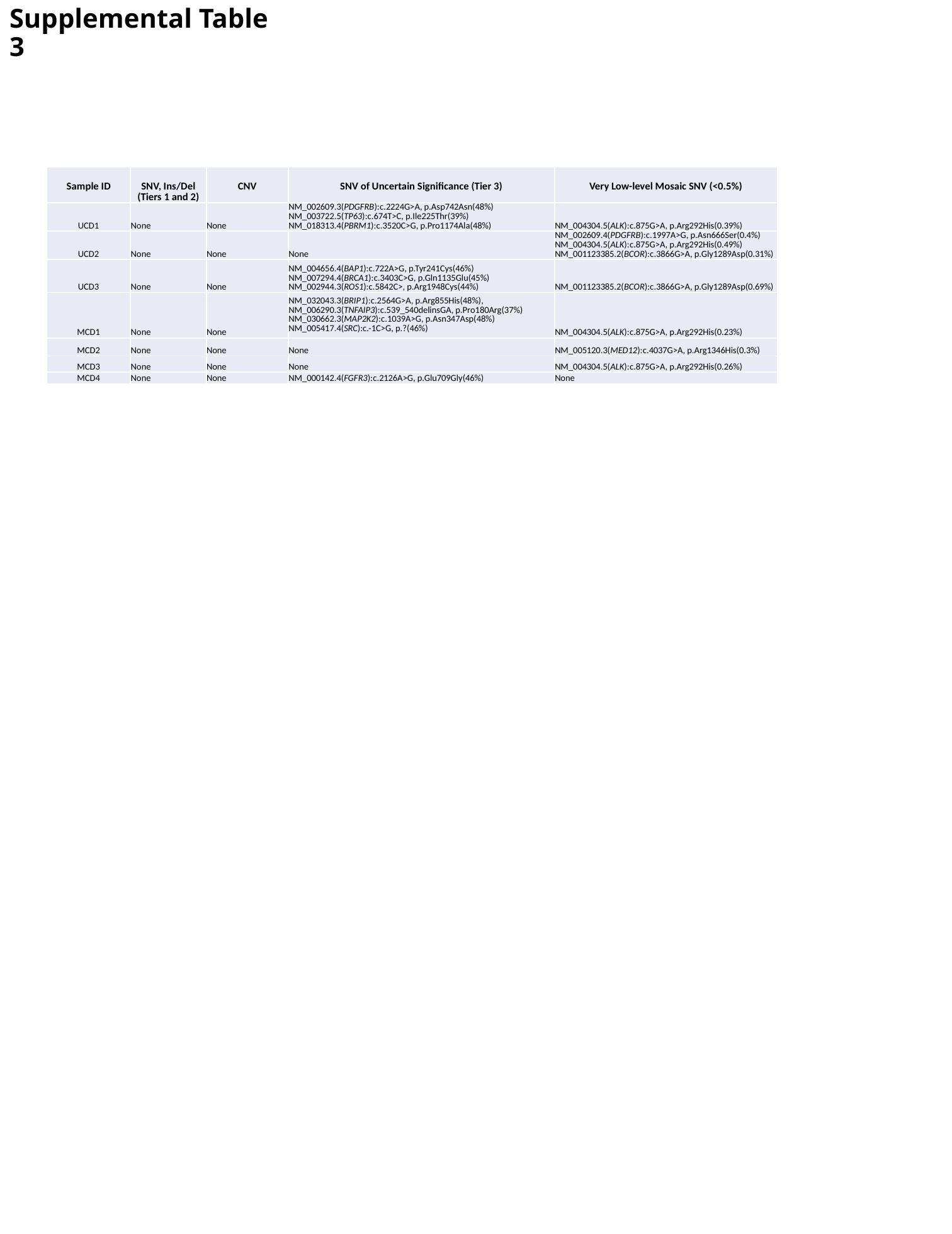

Supplemental Table 3
| Sample ID | SNV, Ins/Del (Tiers 1 and 2) | CNV | SNV of Uncertain Significance (Tier 3) | Very Low-level Mosaic SNV (<0.5%) |
| --- | --- | --- | --- | --- |
| UCD1 | None | None | NM\_002609.3(PDGFRB):c.2224G>A, p.Asp742Asn(48%) NM\_003722.5(TP63):c.674T>C, p.Ile225Thr(39%) NM\_018313.4(PBRM1):c.3520C>G, p.Pro1174Ala(48%) | NM\_004304.5(ALK):c.875G>A, p.Arg292His(0.39%) |
| UCD2 | None | None | None | NM\_002609.4(PDGFRB):c.1997A>G, p.Asn666Ser(0.4%) NM\_004304.5(ALK):c.875G>A, p.Arg292His(0.49%) NM\_001123385.2(BCOR):c.3866G>A, p.Gly1289Asp(0.31%) |
| UCD3 | None | None | NM\_004656.4(BAP1):c.722A>G, p.Tyr241Cys(46%) NM\_007294.4(BRCA1):c.3403C>G, p.Gln1135Glu(45%) NM\_002944.3(ROS1):c.5842C>, p.Arg1948Cys(44%) | NM\_001123385.2(BCOR):c.3866G>A, p.Gly1289Asp(0.69%) |
| MCD1 | None | None | NM\_032043.3(BRIP1):c.2564G>A, p.Arg855His(48%), NM\_006290.3(TNFAIP3):c.539\_540delinsGA, p.Pro180Arg(37%) NM\_030662.3(MAP2K2):c.1039A>G, p.Asn347Asp(48%) NM\_005417.4(SRC):c.-1C>G, p.?(46%) | NM\_004304.5(ALK):c.875G>A, p.Arg292His(0.23%) |
| MCD2 | None | None | None | NM\_005120.3(MED12):c.4037G>A, p.Arg1346His(0.3%) |
| MCD3 | None | None | None | NM\_004304.5(ALK):c.875G>A, p.Arg292His(0.26%) |
| MCD4 | None | None | NM\_000142.4(FGFR3):c.2126A>G, p.Glu709Gly(46%) | None |
