## Supplemental Figures for "Spatial and Single Cell Mapping of Castleman Disease Reveals Key Stromal Cell Types and Cytokine Pathways"

Supplemental Figure 1.

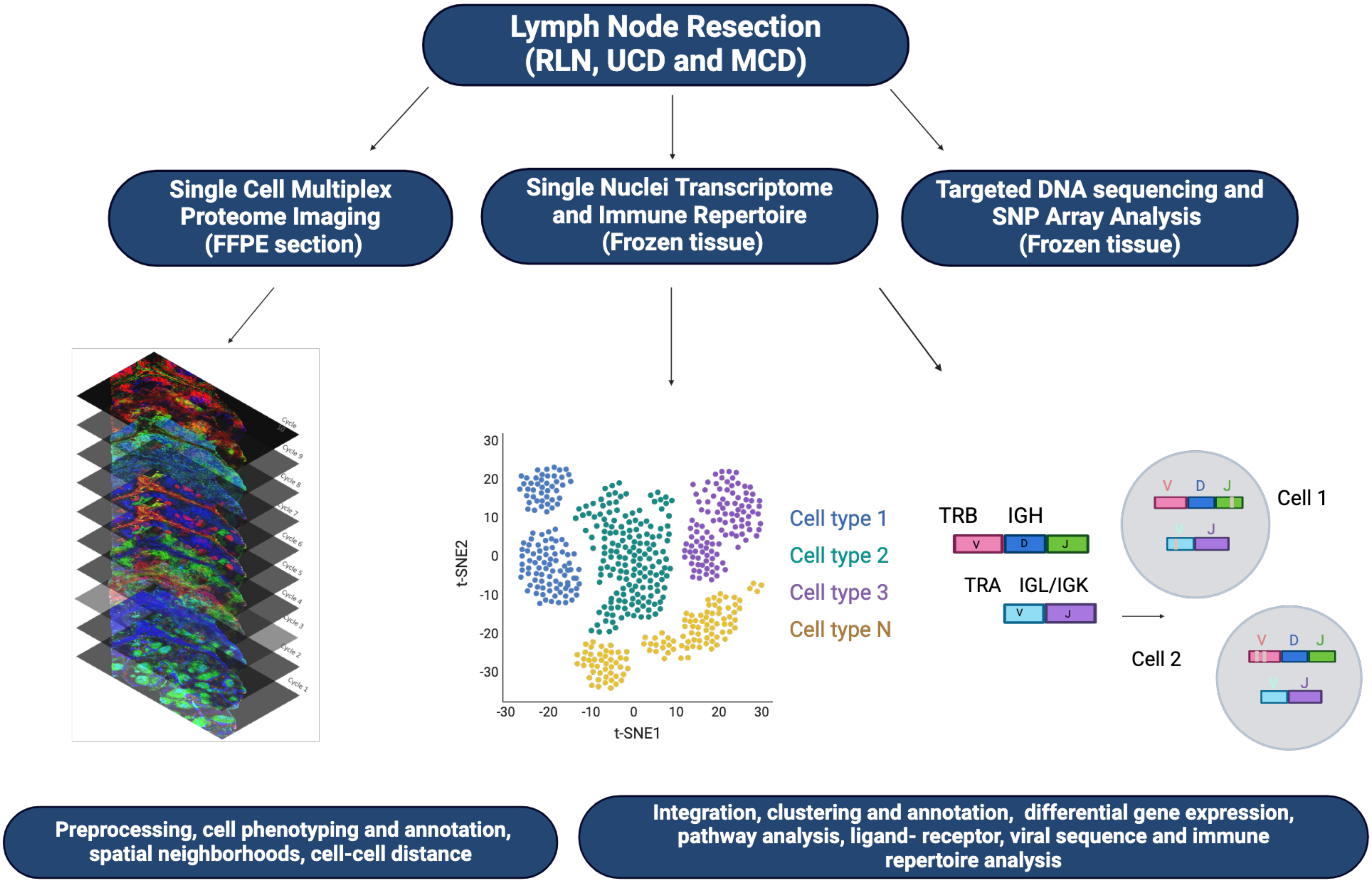

Supplemental figure 2

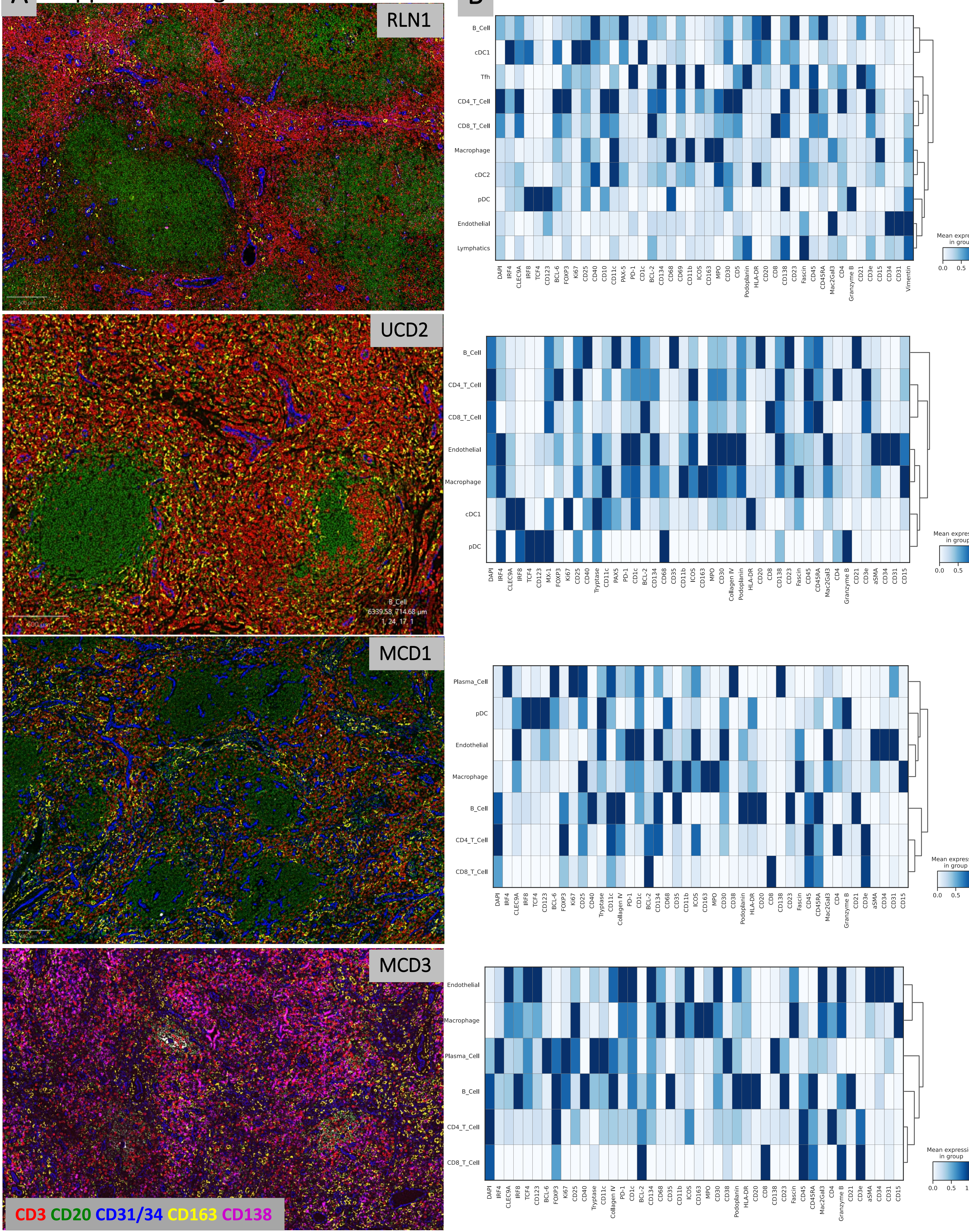

Supplemental Figure 3

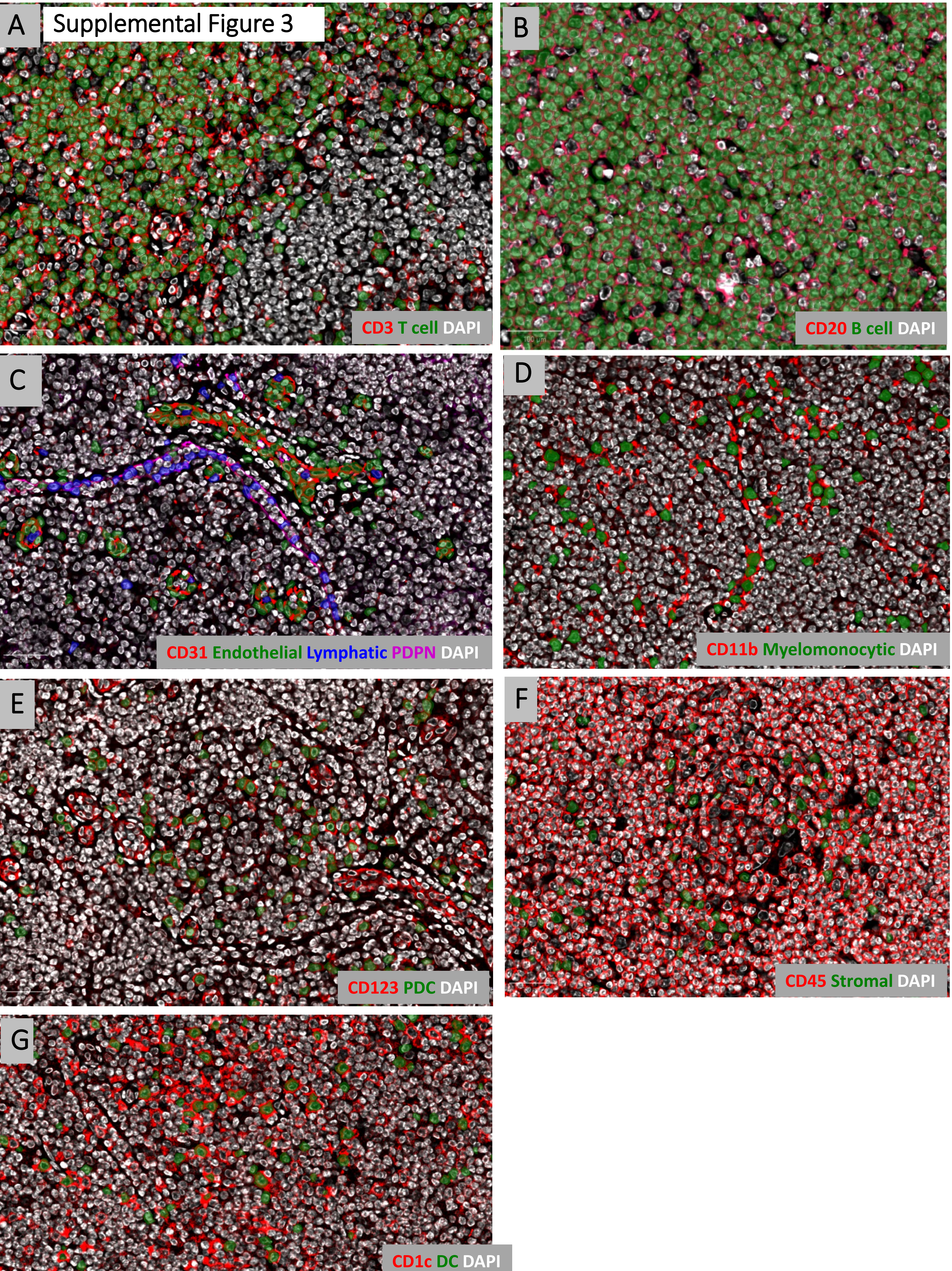

Supplemental Figure 4

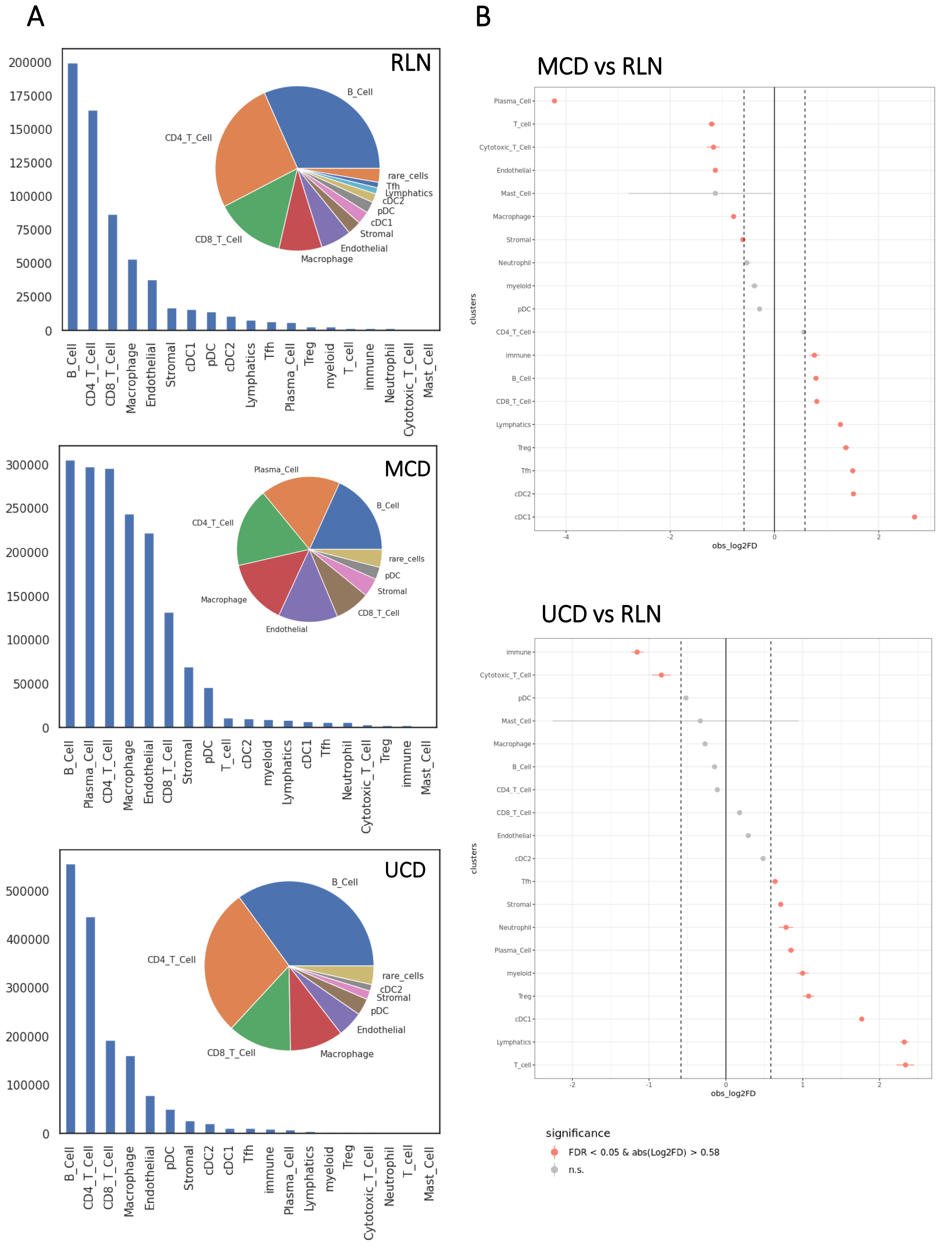

### Supplemental Figure 5

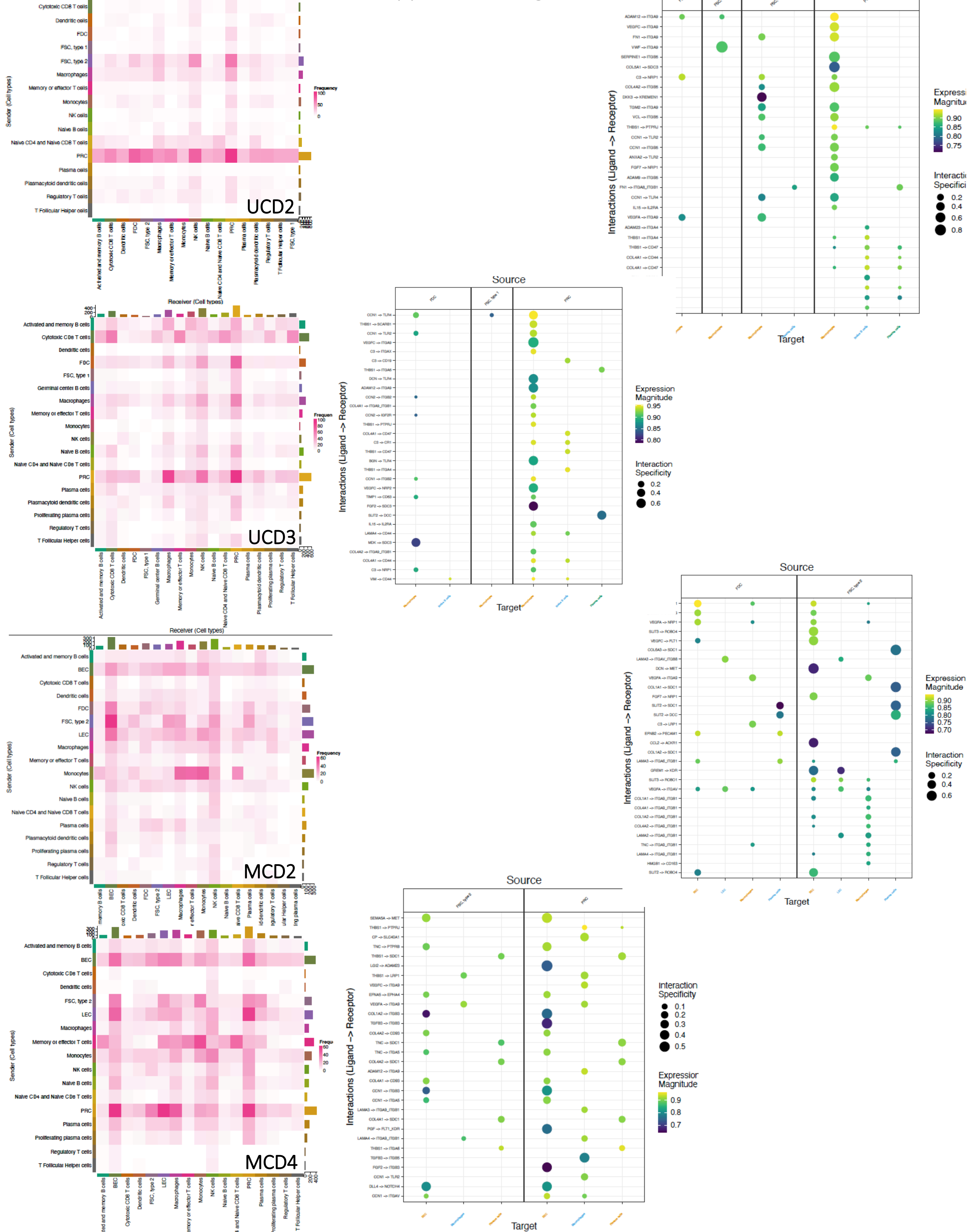

Supplemental Figure 6

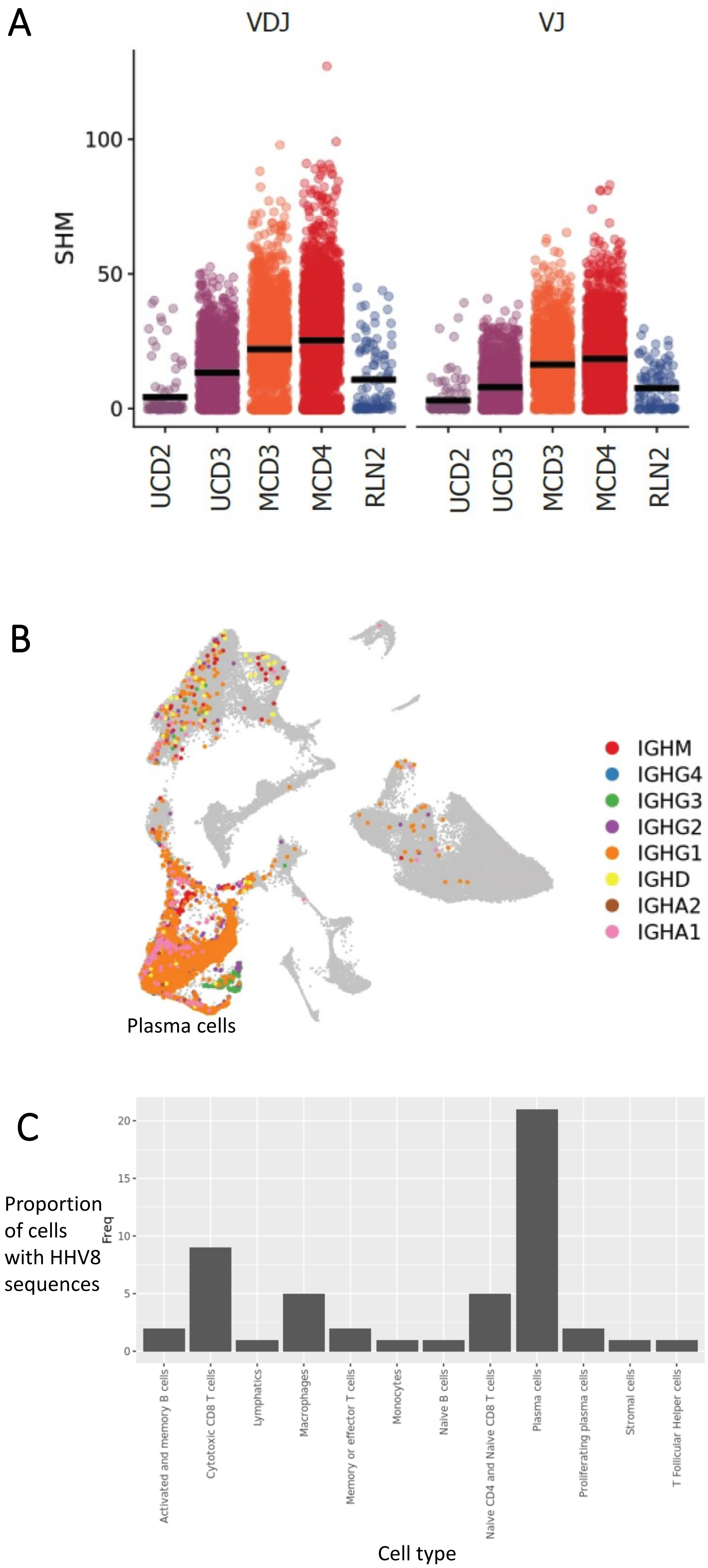
